# A Versatile Enhanced Freeze-Substitution Protocol for Volume Electron Microscopy

**DOI:** 10.1101/2022.05.05.490767

**Authors:** Sébastien Bélanger, Heather Berensmann, Valentina Baena, Keith Duncan, Blake C. Meyers, Kedar Narayan, Kirk J Czymmek

## Abstract

Volume electron microscopy, a powerful approach to generate large three-dimensional cell and tissue volumes at electron microscopy resolutions, is rapidly becoming a routine tool for understanding fundamental and applied biological questions. One of the enabling factors for its adoption has been the development of conventional fixation protocols with improved heavy metal staining. However, freeze-substitution with organic solvent-based fixation and staining has not realized the same level of benefit. Here, we report a straightforward approach including 2% osmium tetroxide, acetone and up to 3% water substitution fluid (compatible with traditional or fast freeze-substitution protocols), warm-up and transition from organic solvent to aqueous 2% OsO_4_. Once fully hydrated, samples were processed in aqueous based potassium ferrocyanide, thiocarbohydrazide, osmium tetroxide, uranyl acetate and lead acetate before resin infiltration and polymerization. We observed a consistent and substantial increase in heavy metal staining across diverse and difficult-to-fix test organisms and tissue types, including plant tissues, nematodes, yeast, and bacteria. Our approach opens new possibilities to combine the benefits of cryo-preservation with enhanced contrast for volume electron microscopy in diverse organisms.

## Introduction

A number of three-dimensional (3D) approaches have been developed that enable intermediate and high-resolution imaging of cells and tissues, each with their own merits and limitations (Watanabe *et al*., 2014; Collman *et al*., 2015; Mahamid *et al*., 2015; Sydor *et al*., 2015; Hoffman *et al*., 2020; Otegui, 2020; Wu *et al*., 2020). Volume electron microscopy (vEM), is particularly suitable when the collection of nanometer scale data from relatively large samples (100s to 1000s of um^3^) and 100s to 1000s of serial sections of resin embedded specimen is required (Titze and Genoud, 2016). This can be achieved by generating consecutive sections arranged in arrays on a substrate, termed array tomography (AT) (Mendenhall, Kuwajima and Harris, 2017; Smith, 2018) or by the serial removal of a thin surface layer and imaging the exposed block-face (Narayan *et al*., 2014; Guérin *et al*., 2019; Lippens *et al*., 2019). With AT, sections cut into ribbons and attached to a surface, post-stained with heavy metals and rendered conductive, allows many traditional fixation/staining protocols as well as affinity probe labeling for correlative microscopy. For serial block-face imaging, an in-chamber ultramicrotome repeatedly shaves the resin surface using a diamond knife and is known as serial block-face scanning electron microscopy (SBF-SEM)(Denk and Horstmann, 2004). Alternatively, a focused ion beam “mills” the resin surface, termed focused ion beam scanning electron microscopy (FIB-SEM)(Narayan and Subramaniam, 2015). Both SBF-SEM or FIB-SEM rely on heavy metal-induced backscatter electrons to generate image contrast, and all metal staining steps of the bulk sample must be performed “*en bloc*,” prior to resin infiltration and polymerization. Furthermore, image quality (signal-to-noise), structure, resolution and sample conductivity are highly dependent upon the levels of metal staining of the sample constituents. Arguably, one of the significant technical advances that helped propel the adoption of SBF-SEM for volume electron microscopy (vEM) studies was the amplification of metal staining via osmium-thiocarbohydrazide-osmium (OTO), often with potassium ferrocyanide, lead and uranium salts (Deerinck *et al*., 2018).

Conventional fixative protocols at ambient temperatures often result in subcellular changes due to fixation artifacts. Immobilizing cellular structures within milliseconds, using freezing, offers optimal morphological preservation of many cellular structures, preserving them in their near-native state (Gilkey and Staehelin, 1986). While freeze-substitution fixation with organic solvent-based staining protocols provides remarkable cellular renditions via post-stained resin sections and transmission electron microscopy (TEM), it often results in very low contrast in certain cell structures (i.e., membranes and cell walls) and overall intense metal staining of traditional freeze-substitution preparations comparable to aqueous OTO remains elusive. Poor membrane visibility often can be addressed by the addition of up to 5% water in the freeze-substitution solution (Buser and Walther, 2008), or other substitution fluid modifications (Guo *et al*., 2020). SBF-SEM in particular is more sensitive to reduced sample conductivity when samples have large non-cellular voids (i.e., vacuoles and/or air spaces). The use of organic solvent-based OTO and/or *en bloc* lead and uranium salts have indeed helped improved contrast and signal-to-noise for vEM (Tsang *et al*., 2018; Czymmek *et al*., 2020), including increased chamber pressure and local gas injection strategies (Deerinck *et al*., 2018) and the inclusion of conductive resins (Nguyen *et al*., 2016) to suppress charge. Despite these improvements, the level of metalization, compared with aqueous OTO protocol counterparts, still limits a number of SBF-SEM and FIB-SEM experiments where improved resolution (x-y, z), speed of acquisition and sample tolerance to beam dosage are required.

To address this, we developed an approach that builds on the work of others using rehydration strategies with HPF freeze-substituted samples (Ripper, Schwarz and Stierhof, 2008; Tsang *et al*., 2018). We transitioned from organic solvent to an aqueous osmium tetroxide solution upon warm-up, and followed this with an aqueous solvent-based metal staining protocol designed to improve uniformity of OTO preparations in larger samples (Hua, Laserstein and Helmstaedter, 2015; Genoud *et al*., 2018; Duncan *et al*., 2022). Our strategy was based upon the notion that the reduced solubility and staining capacity of metal stain components in polar organic vs aqueous solvents for cryo-preparations could be surmounted and further improved. We developed and evaluated our <u>F</u>reeze-<u>S</u>ubstitution and <u>aq</u>ueous <u>OTO</u> protocol (FSaqOTO) using several diverse and challenging plant, animal, and yeast test samples. We included side-by-side aqueous vs organic FS comparisons that demonstrated substantial signal improvement and distinctive staining patterns of certain cell structures with the FSaqOTO protocol. Additionally, our FSaqOTO protocol was versatile; it is compatible with manual, automated and quick-freeze substitution warm-up regimes. While not a panacea, our current FSaqOTO protocol is likely suitable for many organisms and studies for which cryo-preservation is the gold-standard approach, but aqueous enhanced metalization is required to substantially improve signal-to-noise, throughput, and enhanced staining of certain cell structures for intermediate resolution vEM structural studies.

## Materials and Methods

### High-Pressure Freezing Approach for Tested Specimen

Plant: Excised root tips and anthers (0.8 mm and 1.5 mm) were excised via scalpel from barley (*Hordeum vulgare ssp. vulgare*) and placed in 3 mm gold-coated copper planchettes (one Type A, Cat # 16770152 and one Type B, Cat# 16770152, cavity space 400 um) with 50 mM sucrose in 75 mM PIPES buffer as a space filler and high-pressure frozen.

Nematode: *Caenorhabditis elegans* were maintained on MYOB agar plates seeded with *E. coli* OP50 bacteria and prepared for high pressure freezing as described previously (Rahman *et al*., 2021). Briefly, nematodes were collected in cellulose capillary tubes (Leica catalog number 16706869) and placed in 3 mm gold-coated copper planchettes (Type A and B, cavity space 300 um) with 20% dextran as a space filler and high-pressure frozen.

Yeast: *Saccharomyces cerevisiae* (strain BJ5494) was grown in YEPD mediim to mid-log phase in a shaker flask at 60 RPM and 32°C, harvested by gentle centrifugation on a tabletop device to create a yeast paste and placed in 3mm gold-coated copper planchettes (Type A and B, cavity space 100 um) without additional space filler and high-pressure frozen.

### Freeze-substitution and Rehydration

All specimens were frozen in a Leica EM ICE High-Pressure Freezer (Leica Microsystems, Inc., Buffalo Grove, IL, USA), processed for freeze-substitution, heavy metal stained and resin infiltrated according to the general protocol overview shown in Figure 1. The overall approach was robust and allowed variations in substitution fluid, resin formulation and warm-up strategy.

**Figure 1.**
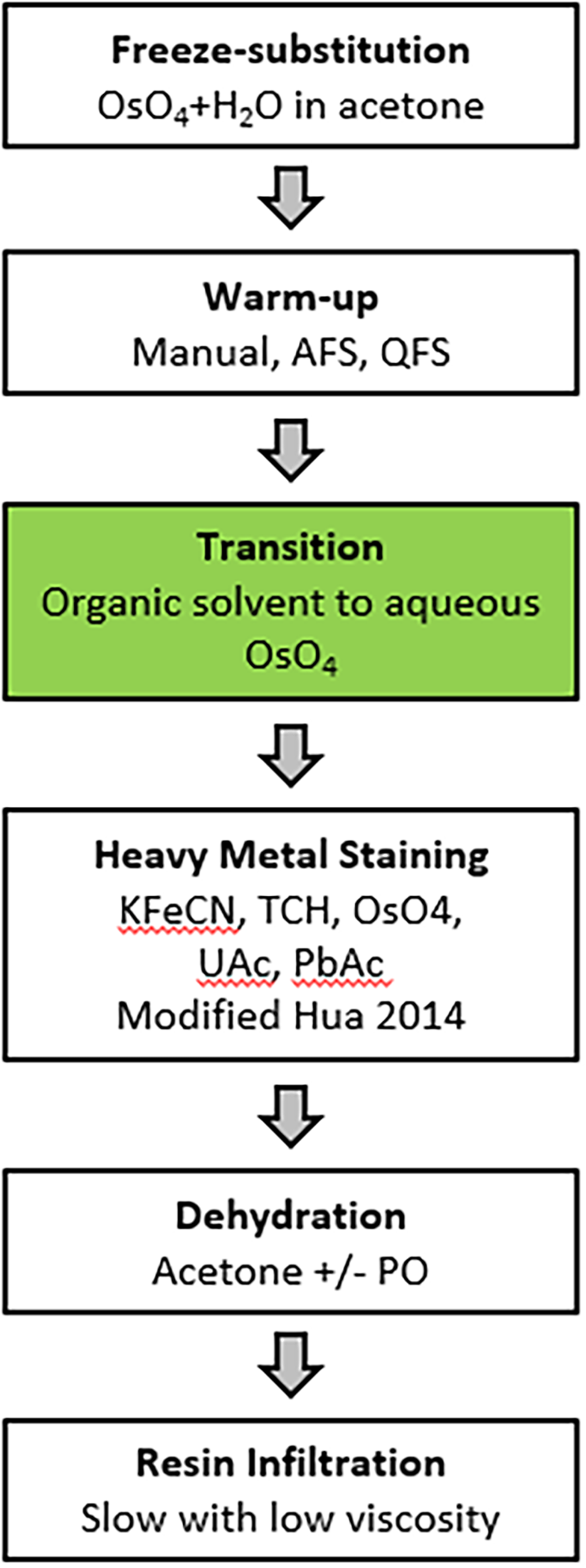
FSaqOTO general protocol overview. The major steps in the FSaqOTO protocol include freeze-substitution, warm-up to 0-4°C, the critical step (green filled) transition from organic solvent to aqueous buffer with osmium tetroxide (OsO_4_), heavy metal staining with potassium ferrocyanide (KFeCN), thiocarbohydrazide (TCH), osmium tetroxide (OsO_4_), uranyl acetate (UAc) and lead acetate (PbAc). Dehydration in acetone with/without propylene oxide and resin infiltration, slow (increased duration and/or steps) and low viscosity.

HPF frozen plant materials were placed in mPrep/s capsules (Microscopy Innovations, Cat # 2250) without mPrep screens and held in mPrep CPD Holder (Microscopy Innovations, Cat # 2250) that was then placed in a 50 ml PTFE beaker (EMS Diasum, Cat# 60942) containing LN_2_ frozen substitution fluid (2% osmium tetroxide in acetone with 3% water) and subsequently moved to -85°C for 3–4 days (Supplementary Data 1). Aluminum planchettes and substitution containers should be avoided as the heavy metal staining steps react with aluminum (i.e. TCH) turning the solution black and subsequent use of aluminum containers with this chemistry resulted in their failure (leakiness). Samples were then transferred and held at -55°C and -20°C for 3 hours each and moved to 4°C for 1 hour and room temperature for 1 hour. At 4°C, a second mPrep/s capsule was inserted into each initial mPrep/s capsule to entrap and immerse the small specimen during subsequent aqueous-based processing steps. This prevented specimen loss and ensured the buoyant plant specimens were completely submerged in all reagents, thus minimizing uneven staining and/or inadequate reagent exchange. The critical organic solvent to aqueous transition step was performed at 4°C, where samples were rehydrated (20 minutes each step) in 1% OsO_4_ using the following graded series of H_2_O:acetone 1:4, 1:1, 4:1 and finally moved to room temperature in 1% OsO_4_ in 0.1 M sodium cacodylate buffer for 1 hour.

For protocol comparison purposes, freeze-substitution was applied using a traditional Automated Freeze-substitution System (AFS, Leica Microsystems) (Rahman *et al*., 2021) without FSAqOTO, and Quick Freeze-Substitution (QFS) (Mcdonald and Webb, 2011) with our enhanced aqueous metalization approach to the model organisms *S. cerevisiae* and *C. elegans*. With AFS, the freeze-substitution cocktail was 1% OsO4 and 1% glutaraldehyde in 1% H_2_O in acetone. 1 mL of freeze substitution cocktail was placed in each cryovial and frozen under liquid nitrogen. The planchettes holding the frozen samples were transferred under liquid nitrogen vapors into the cryovials containing the frozen freeze substitution cocktail. The cryovials were then capped and placed in the Leica AFS 2 chamber (set to -90°C) in aluminum containers with pre-cooled ethanol. The AFS program was run as follows: -90°C for 48 hours, - 90°C to -60°C for 6 hours (5°C/hour), -60°C for 2 hours, -60°C to -20°C for 8 hours (5°C/hour), - 20°C for 2 hours, -20°C to 0°C for 4 hours (5°C/hour). After freeze substitution, the samples were rinsed with pre-cooled (to 0°C) acetone 3 times for 10 minutes each. After the third wash, samples were removed from the AFS chamber and allowed to reach room temperature. Samples were placed in 1% uranyl acetate in acetone for 1 hour at room temperature followed by 3 acetone washes for 10 minutes each. Resin infiltration (using Polybed 812, DDSA, NMA, Polysciences, Inc. BDMA, Electron Microscopy Sciences) was done at room temperature as follows: *C. elegans* - 1:3 resin: acetone for 2 hrs, 1:1 resin: acetone for 2 hrs, 3:1 resin: acetone overnight, 100% resin for 5 hours. *S. cerevisiae* - 1:3 resin: acetone for 2 hrs, 1:1 resin: acetone overnight, 3:1 resin: acetone for 3 hours, 100% resin overnight. Samples were embedded in molds with fresh resin and polymerized in a 60°C oven for 2 days.

In addition to AFS, we applied a Quick Freeze Substitution aqueous OTO protocol for both the *S. cerevisiae* and *C. elegans* samples. Briefly, after high pressure freezing in an Leica EM ICE, the FSaqOTO protocol was performed largely following Rahman (Rahman *et al*., 2021) and McDonald (McDonald, 2014), but with slight changes to the actual freeze substitution solvent and stain mixtures (referred to as SubMix). For yeast samples, a SubMix of 1% w/v OsO4 and 1% w/v UA dissolved in 90% acetone + 5% methanol + 5% water was prepared. For *C. elegans*, a SubMix of 1% w/v OsO4 dissolved in 97% acetone + 3% water was prepared. Briefly, 1 ml of these SubMixes were aliquoted into cryovials and inserted into a metal block, which was laid on its side in an ice bucket. The bucket was filled with liquid nitrogen, and after the SubMixes were frozen, the high-pressure frozen planchettes with the samples were quickly transferred to the cryovials. The liquid nitrogen was poured out and dry ice packed in (for a detailed step-by-step protocol with images and catalog numbers, please see Rahman (Rahman *et al*., 2021). The ice bucket was put on a rotary shaker set at 60 rpm, and rotated for 3 hours with the lid on, 1 hour with the dry ice poured out but the lid still on, and finally with the lid off until the temperature reached 0°C, typically 1 hour. At this point, rather than continue with the freeze substitution in the mostly anhydrous SubMix, the metal block was surrounded by ice packs to maintain temperatures around 0°C. Then, the SubMix was carefully replaced with previously made and cooled dilutions of the SubMix. Thus, the samples were incubated with 1 mL pre-chilled solutions of SubMix:water ratios of 3:1, 1:1, 1:3 each for 20 minutes, with the shaker still set at 60 rpm. Finally, the solution was replaced with 1% OsO4 in 0.1 M sodium cacodylate buffer and allowed to equilibrate to room temperature for 1 hour. After this step, the Hua heavy metal staining protocol (Hua, Laserstein and Helmstaedter, 2015) was largely followed, except we used PolyBed resin, hard formulation.

The resin embedded samples were trimmed and sectioned to expose the biosample on the section face, after which the resin block was cut, mounted on a stub, gold coated and introduced into a CrossBeam550 (ZEISS, Oberkochen, Germany) for high-resolution FIB milling and SEM imaging.

### Heavy Metal Staining

We subsequently performed a modified version of the OTO staining method developed by Hua *et al*. (2015) for sample metallization of large mouse brain specimens. We removed the osmium solution and placed directly into 2.5% potassium ferrocyanide (#9387-100G; Sigma-Aldrich, St Louis, MO, USA) in 0.1 M sodium cacodylate at pH 7.4 for 90 minutes at room temperature. We then washed samples (2 × 30 minutes in H_2_O) and transferred samples into 1% TCH (#21900; EMS, Hatfield, PA, USA) in H_2_O for 45 minutes at 40°C. We washed samples (2 × 30 minutes in H_2_O) at room temperature again and transferred samples into 2% OsO_4_ in H_2_O for 90 minutes at room temperature. We washed samples (2 × 30 minutes in H_2_O) and transferred samples in an unbuffered 1% aqueous uranyl acetate at 4°C, overnight. The samples in 1% aqueous uranyl acetate were then moved to 50°C for 2 hours and washed (2 × 30 minutes in H_2_O) and transferred to lead aspartate in H_2_O for 2 hours at 50°C. The lead aspartate was prepared with 0.04 g L-aspartic acid (#a9256; Sigma-Aldrich, St Louis, MO, USA) and 0.066 g lead nitrate (#203580-10G; Sigma-Aldrich, St Louis, MO, USA) in 10 ml water and the pH was adjusted to 5.5. Finally, samples were washed (2 × 30 minutes in H_2_O) at room temperature.

### Sample Dehydration, Infiltration and Embedding

Prior to infiltration and sample embedment, we performed a graded dehydration series of 30%, 50%, 70%, 80%, 90%, 100% and 100% acetone at 4°C for 30 min each. The dehydrated samples were exchanged 2X in 100% of propylene oxide (#20401; EMS, Hatfield, PA, USA) for 30 min each. Samples were then resin infiltrated with a graded series of 25%, 50%, 75% and 100% Quetol resin (hard formulation) in propylene oxide without DMP-30 (#14640; EMS, Hatfield, PA, USA) at room temperature for 24 hours each step on a rocking platform to enhance resin infiltration. All samples were removed from the mPrep/s capsules at the 50% acetone graded series step and then processed in glass vials for remaining steps. Subsequently, two overnight 100% resin exchanges were made with DMP-30. Finally, samples were embedded in freshly made 100% Quetol using flat embedding molds (#70900; EMS, Hatfield, PA, USA).

All major steps in sample preparation conditions from fixation to resin are detailed in Supplementary Data 2.

### SBF-SEM, FIB-SEM, STEM and TEM Image Acquisition

Embedded tissues were mounted on 1.4 mm standard Gatan flat pin using silver conductive epoxy (Chemtronics, CW2400, Kennesaw, GA, USA), trimmed and sectioned using the Leica Ultracut UCT (Leica Microsystems Inc., Buffalo Grove, IL, USA). Due to the extremely opaque nature of the metalized samples prepared in this way, sample quality was assessed any of the following approaches: whole block imaging by X-ray microscopy, unstained semi thick sections (350 nm) via light microscopy, or ultrathin sections (∼70 nm) by transmission electron microscopy. For SBF-SEM, pins with resin blocks were sputter coated with an ∼50 nm layer of gold/palladium. Then, over 1000 sequential images at a 10k × 10k pixel resolution and 5nm x-y, 50nm z-step size were collected on a ZEISS GeminiSEM 300 SEM at 5kV (current of 1 pA, and 3 µs dwell time) using a Gatan 3-View® 2XP and local N_2_ gas injection via a Focal Charge Compensation (FCC) needle set between 10-35%.

FIB-SEM imaging was performed on a Zeiss CrossBeam 550, using the ATLAS3D module, as previously described (Narayan *et al*., 2014). After the specimen were coated with a patterned platinum and carbon pad, images were acquired at either 3 or 5 nm pixel sampling and 4 µs total pixel dwell time, with electron beam parameters of 1.5 kV and 1 nA and the grid voltage at the in-column backscatter detector set at 900 V. The FIB-SEM was operated at 30 kV, 1.5 µA, with a step size of either 9 nm or 15 nm, respectively. The resulting image stacks were registered, contrast inverted and binned to yield isotropic image volumes. STEM images were acquired on a Zeiss GeminiSEM 450 equipped with a STEM detector and operated at 30kV. Images were acquired at 3, 10, or 15 nm x-y pixel sampling.

All manuscript figures and corresponding imaging conditions are detailed in Supplementary Data 3.

### Image Processing, Segmentation, Reconstruction and Visualization of Specimen Volumes

Three-dimensional volumes of selected specimens using the Object Research Solutions (ORS, Montréal, Canada) Dragonfly Version 2020.2 Build 941. For segmentation and deep learning, an ∼180 image 10k x10k subset of the full stack was processed using the ORS Segmentation Wizard. Three slices were fully trained representing eight target features. A three-level Sensor 3D deep learning model with a patch size of 64 (Supplementary Data 4) yielded good results for our aqueous OTO freeze-substitution protocol for plant cell walls, mitochondria, Golgi and vesicles in 3D surface renderings (Fig. 1E).

## Results

### FSaqOTO Protocol Enhanced Staining of Whole-Mount Plant Tissues

In plants, some of the primary challenges when applying vEM is related to the cell walls, large air voids and water filled vacuoles which impede not only fixation, staining, but also impair overall sample conductivity, resulting in sensitivity to beam damage and charging artifacts. These difficulties are further exacerbated when working with cryo-fixation and freeze-substitution protocols, as organic solvent-based metal staining can be relatively limited due to low solubility, reduced chemical reactivity and overall poor staining of certain cell membranes (i.e. endoplasmic reticulum, nuclear envelope, mitochondria and thylakoids). Recent work applying vEM to freeze-substituted anthers with organic solvent based OTO *en bloc* staining in combination with SBF-SEM and FCC, did improve accessibility for many biological questions in these important plant structures in our hands (Duncan *et al*., 2022). Despite considerable effort, we were unable to realize the full contrast benefits observed in traditional conventionally fixed specimens using aqueous-based OTO protocols while maintaining a strictly organic solvent-based processing routine after cryo-fixation. However, based on our recent success using conventional fixation with tobacco leaf tissues, the Hua protocol (Hua, Laserstein and Helmstaedter, 2015) was shown to be highly suitable for vEM and X-Ray microscopy. Thus, we reasoned that a modification of our freeze-substitution protocol with a graded transition from an organic solvent-based (in our case acetone) 1% OsO_4_ solution (containing H_2_O), to full rehydration in 1% OsO_4_ aqueous buffer, followed by the Hua heavy metal staining steps, would enhance staining of freeze-substituted plant samples. Indeed, this concept worked with good effect in barley roots (Fig. 2) and anthers (Fig. 3) with SBF-SEM and FCC. A low magnification cross-section of a HPF FSaqOTO prepared barely root near the meristem (Fig. 2A) showed high-density ground cytoplasmic matrix of the epidermal and cortex cells and elevated staining and electron density of the plant cell walls and vacuolar compartments (Fig. 2B-D), Golgi apparatus and trans Golgi secretory vesicles (Fig. 2C). Additionally, the plasma membrane, mitochondrial membranes and endoplasmic reticulum, while not heavily stained compared to conventional fixation OTO protocols, were readily discerned (Fig. 2D). Indeed, this aided the success of deep learning segmentation (Sensor 3D, ORS Dragonfly) for 3D renderings (Fig. 2E) of mitochondria (blue), Golgi and secretory vesicles (green) and cell wall (magenta) as well as intersecting plasmodesmata voids (arrows). Interestingly, plasmodesmata, which normally are more electron dense relative to the cell wall via electron microscopy, had inverted contrast (appeared electron transparent in a more electron dense cell wall) (Fig. 2D, Supplementary Data Video_1).

**Figure 2.**
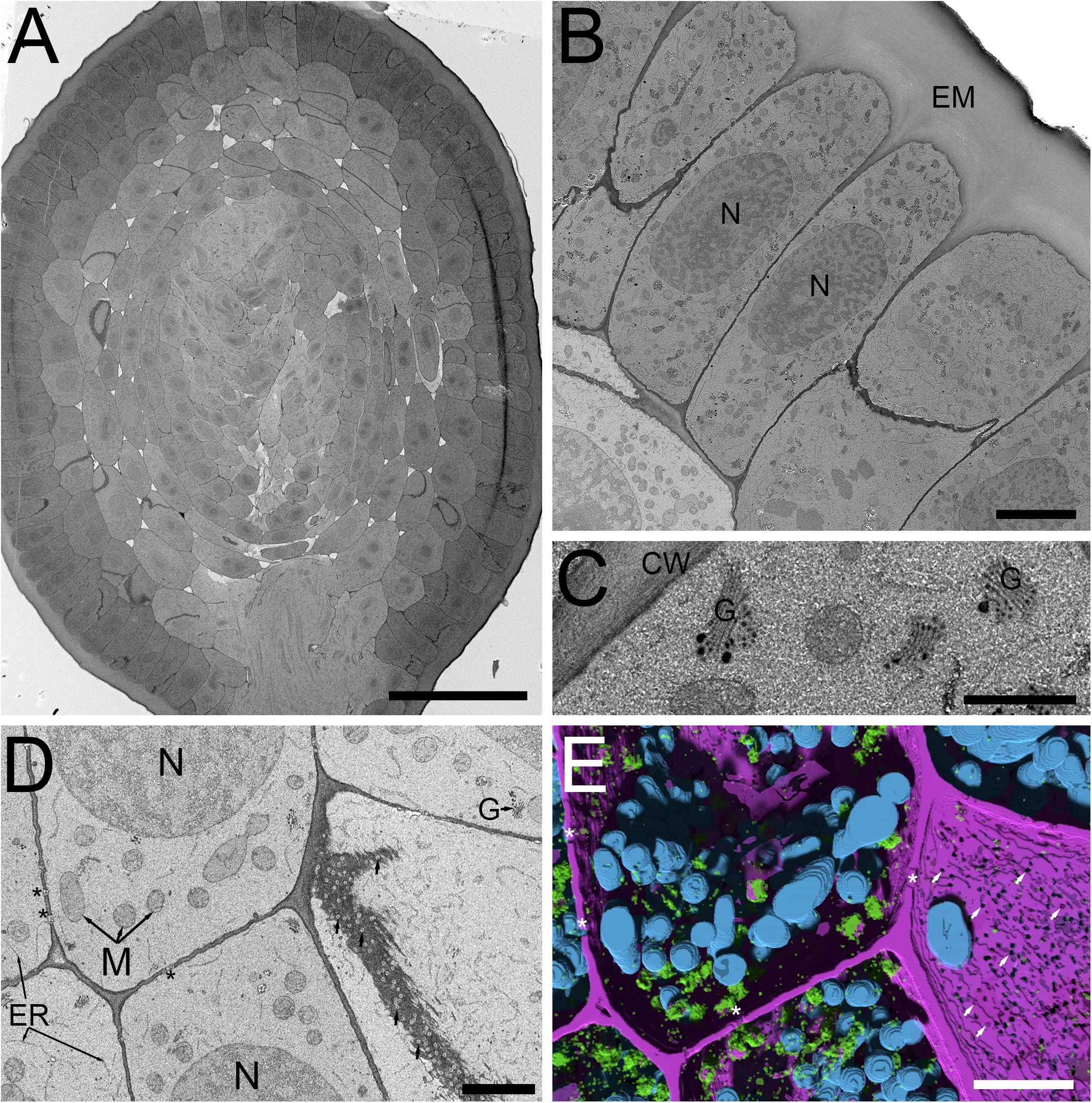
SBF-SEM of high-pressure frozen barley root prepared by FSaqOTO. **A**. SBF-SEM overview of root cross-section near the root apical meristem prepared by FSaqOTO. Scale = 100 µm. **B**. High-resolution SBF-SEM slice from **A** showing the outermost protoderm cells with nuclei (N), a normal array of other organelles and covered by a thin electron dense cell wall with a thick extracellular matrix (EM) on the root exterior. Scale = 5 µm. **C**. High magnification of a protoderm cell with well-stained Golgi membranes, cell wall and trans-Golgi vesicle contents. Scale = 0.5 µm. **D**. Rapidly dividing cells in the ground meristem exhibited well contrasted cell walls (CW) Golgi/vesicles (G) with clearly delineated nuclei (N), mitochondria (M) and electron transparent plasmodesmata (arrows). Scale = 1 µm. **E**. 3D rendering of **D** using Sensor 3D deep learning segmentation for mitochondria (cyan), Golgi and secretory vesicles (green) and cell wall (magenta) as well as intersecting plasmodesmata voids (asterisks). Scale = 1 µm.

**Figure 3.**
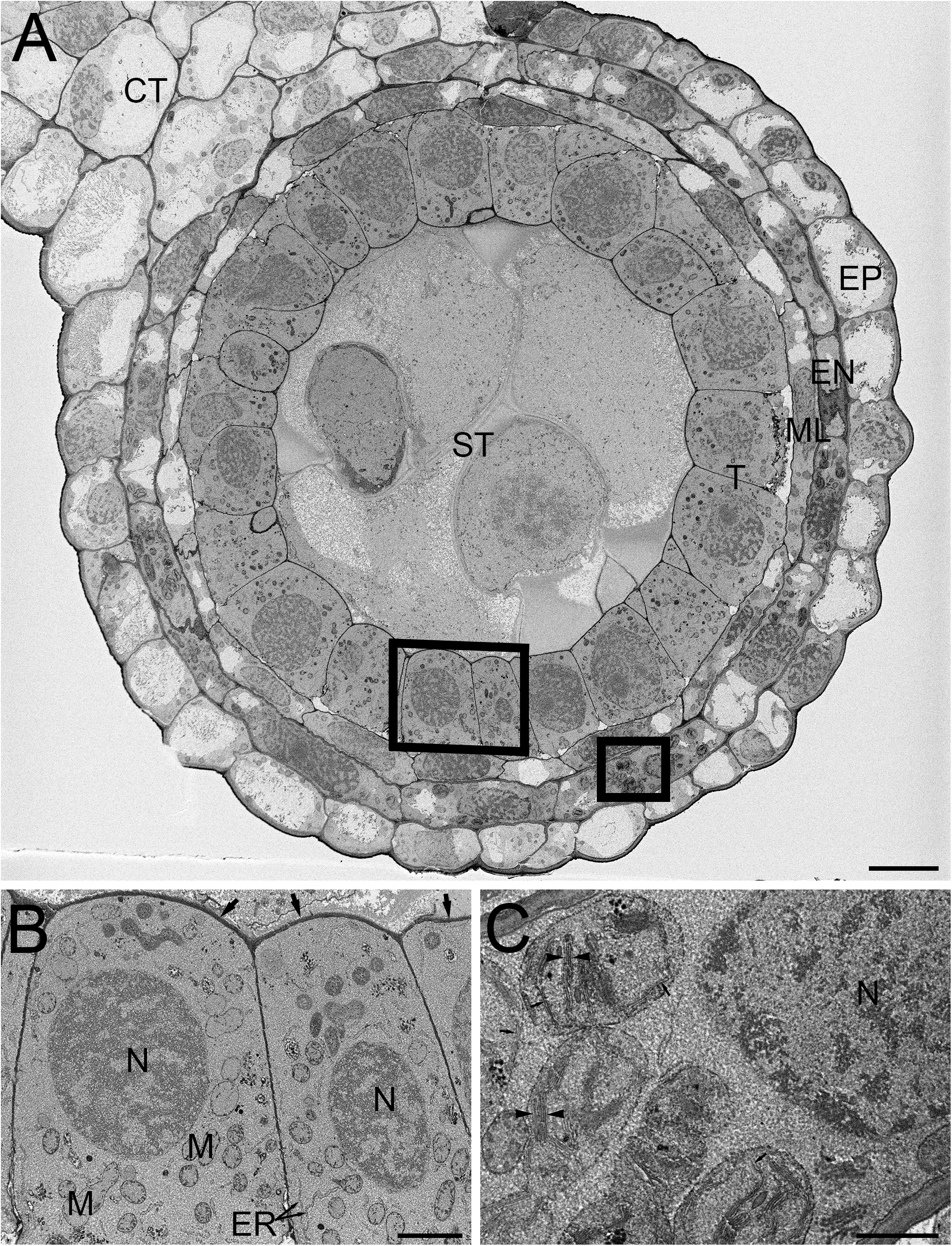
SBF-SEM of high-pressure frozen barley anther prepared by FSaqOTO. **A**. SBF-SEM overview of anther cross-section showing epidermal (EP), endothecium (EN), middle layer (ML), tapetum (T), sporogenous (ST) and connective tissue (CT). Scale = 10 µm. **B**. High magnification slice from region **A** (left box) showing the tapetum cells with nuclei (N), mitochondria (M) endoplasmic reticulum (ER) and covered by a thin electron dense cell wall (arrows). Scale = 1 µm. **C**. High magnification from region **A** (right box) well-stained well-stained chloroplastic membranes (arrows) and grana stacking readily visible (arrowheads). Scale = 0.5 µm.

We then evaluated the FSaqOTO protocol for SBF-SEM and FCC on barley anthers, the male reproductive structure responsible for pollen production in plants. A low magnification overview of a single barley anther lobe allowed a clear assessment of the epidermal (EP), endothecium (EN), middle layer (ML), tapetum (T), sporogenous (ST) and connective tissue (CT) (Fig. 3A). Similar to root tissues (Fig. 2), the FSaqOTO showed elevated electron density in cells with high cytoplasmic density (Fig. 3A-C) and highly contrasted cell walls. Closer inspection of the tapetum (Fig. 3B) revealed good contrast and discrimination of mitochondrial membranes, Golgi apparatus, endoplasmic reticulum and cell wall. While the nuclear envelope was visible, the elevated staining of the heterochromatin and nucleoplasm relative to the nuclear envelope, made it less conspicuous, especially in oblique profiles (Fig. 2D, 3B & C). In the endothecium, chloroplast thylakoid and grana membranes were readily contrasted and resolved (Fig. 3C). Interestingly, in both plant tissues, non-membranous components of the cytoplasm, chloroplast stroma and nucleoplasm appeared to have a granular or textured appearance. In both tissue types, over 1000 consecutive 10,000 × 10,000 pixel serial images were readily obtained at 5 nm x-y pixel resolution and 50 nm z-slice interval.

### *FSaqOTO Protocol Enhanced Staining of Whole-Mount* C. elegans

After verifying improved conductivity and overall elevated staining characteristics of our FSaqOTO protocol using SBF-SEM on challenging plant tissues, we tested if the method also provided similar improved staining with other phylogenetically distinct organisms. Furthermore, we wanted to compare the relative increase and differences in staining of cell structures when a quick freeze substitution (QFS) protocol was used versus a traditional AFS freeze-substitution protocol. QFS is an excellent option for freeze-substitution in the absence of expensive instrumentation, with the added advantage of speed compared to AFS or freezer-based FS methods. For this work, we chose the model organisms *C. elegans* and *S. cerevisiae*; both represented challenging tests for EM staining protocols on account of their cuticle and cell wall, respectively, and were likely to show significant differences in staining with increased metallization, including vEM. For *C. elegans*, we noted a substantial contrast improvement across the organism when side-by-side AFS versus FSaqOTO comparisons of worm longitudinal sections were made under identical image acquisition conditions (Fig. 4A & B). We observed some freezing artifacts of the embryos (Fig. 4A asterisk), which is unsurprising given the large size of the intact worm and chitinaceous shell of the embryo. It is likely that freezing was slowest at these depths, significantly away from the freezing surfaces during the HPF step. Nevertheless, at increased magnification, FSaqOTO of *C. elegans* demonstrated high contrast of various tissue and cell features. The fibers of smooth muscle were clearly delineated, and membranes of spermatheca, including the plasma membrane, mitochondria, and membranous organelles were well stained (Fig. 4C). Similar to plant tissues, these samples had a textured cytoplasm and nucleoplasm. Under matched image acquisition conditions (e.g. Fig. 4C = Fig. 4D), but rescaled display histogram (Fig. 4D only), the traditional AFS protocol lacked strong cytoplasmic and membrane staining, especially for the mitochondria and endoplasmic reticulum.

**Figure 4.**
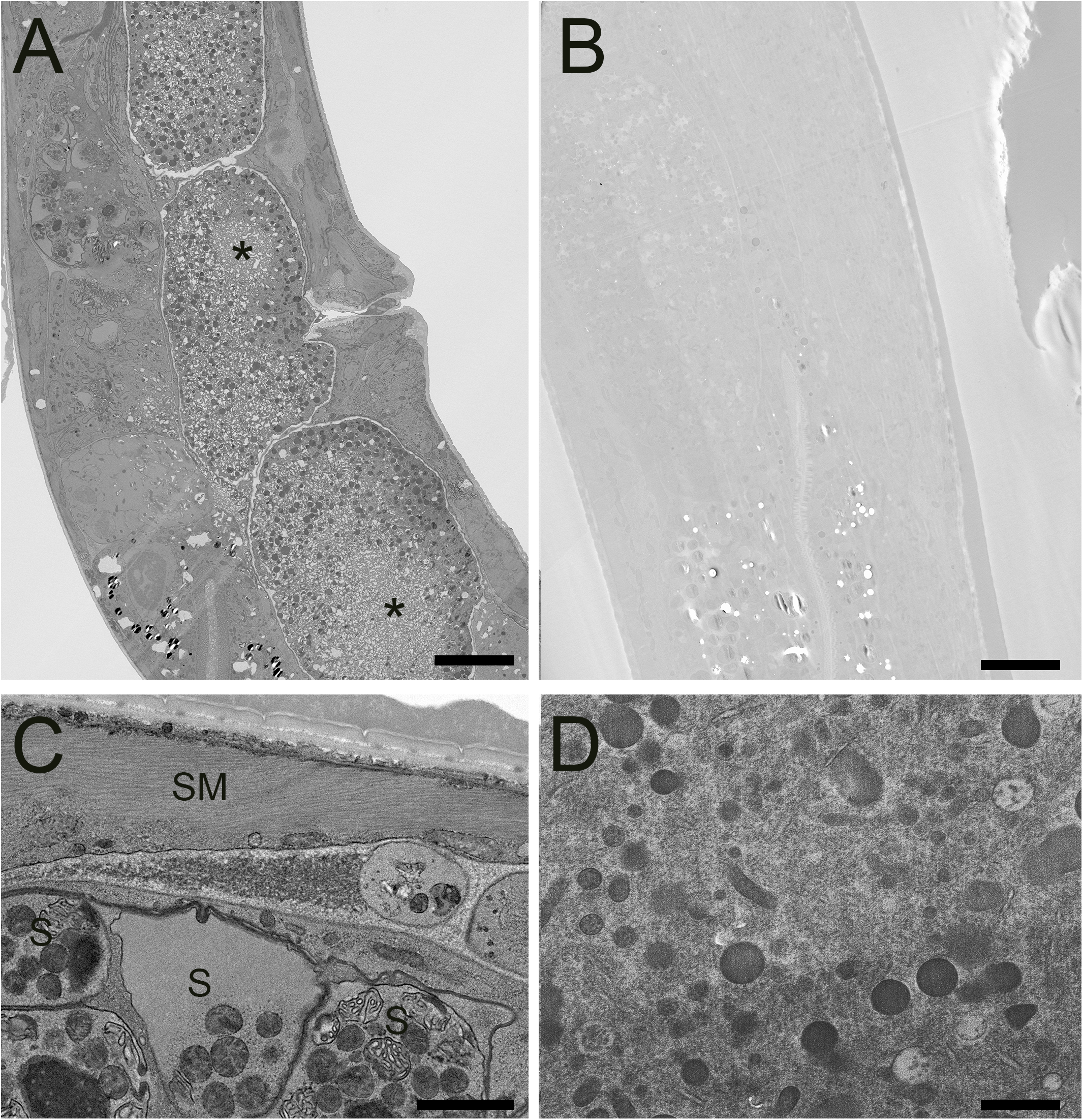
STEM of high-pressure frozen *C. elegans* prepared by FSaqOTO and traditional AFS. **A**. Low resolution overview longitudinal section of the nematode demonstrated very high contrast staining using FAaqOTO and using <u>identica</u>l imaging condition for **B** compared to a traditional AFS staining protocol. Local freeze-damage (asterisk). A & B, scale = 10 µm. **C**. FSaqOTO prepped sample at increased magnification with smooth muscle fibers (SM) clearly delineated, and well-stained membranes of spermathecal (S), including the plasma membrane, mitochondria, other membranous organelles. Scale = 1 µm. **D**. Corresponding high magnification (compare to **4C**) with identical acquisition conditions of an AFS processed *C. elegans* <u>but</u> with a rescaled histogram demonstrated the normal complement of organelles but with notably relatively much reduced contrast, especially most cell membranes. Scale = 1 µm.

Like *C. elegans*, a direct comparison of traditional AFS and FSaqOTO protocols with the yeast, *S. cerevisiae*, evaluated by TEM demonstrated a substantial improvement in staining and contrast for FSaqOTO (Fig. 5A versus 5B) under identical acquisition and image display conditions. Of note is the strong contrast of the yeast cell wall, glycogen, plasma membrane, cisternae (Golgi equivalent) and vacuolar membrane and its contents (Fig. 5A & 5C) compared to its AFS counterpart (Fig. 5B & 5D). As with plant roots, anthers and *C elegans*, the ground cytoplasmic and nucleoplasm matrices exhibited a textured or slightly granular appearance (Fig. 5C). While mitochondria and nuclei staining were notably elevated and visible in FSaqOTO (Fig. 5C & inset) versus traditional AFS prepared yeast, the yeast mitochondrial membranes, nuclear envelope, and endoplasmic reticulum were essentially lacking in the AFS protocol, even after histogram rescaling the identically acquired image (Fig. 5D & inset). FIB-SEM serial-sections of the FSaqOTO sample blocks were consistent with TEM data in contrasted structures (Supplementary Data Video 2).

**Figure 5.**
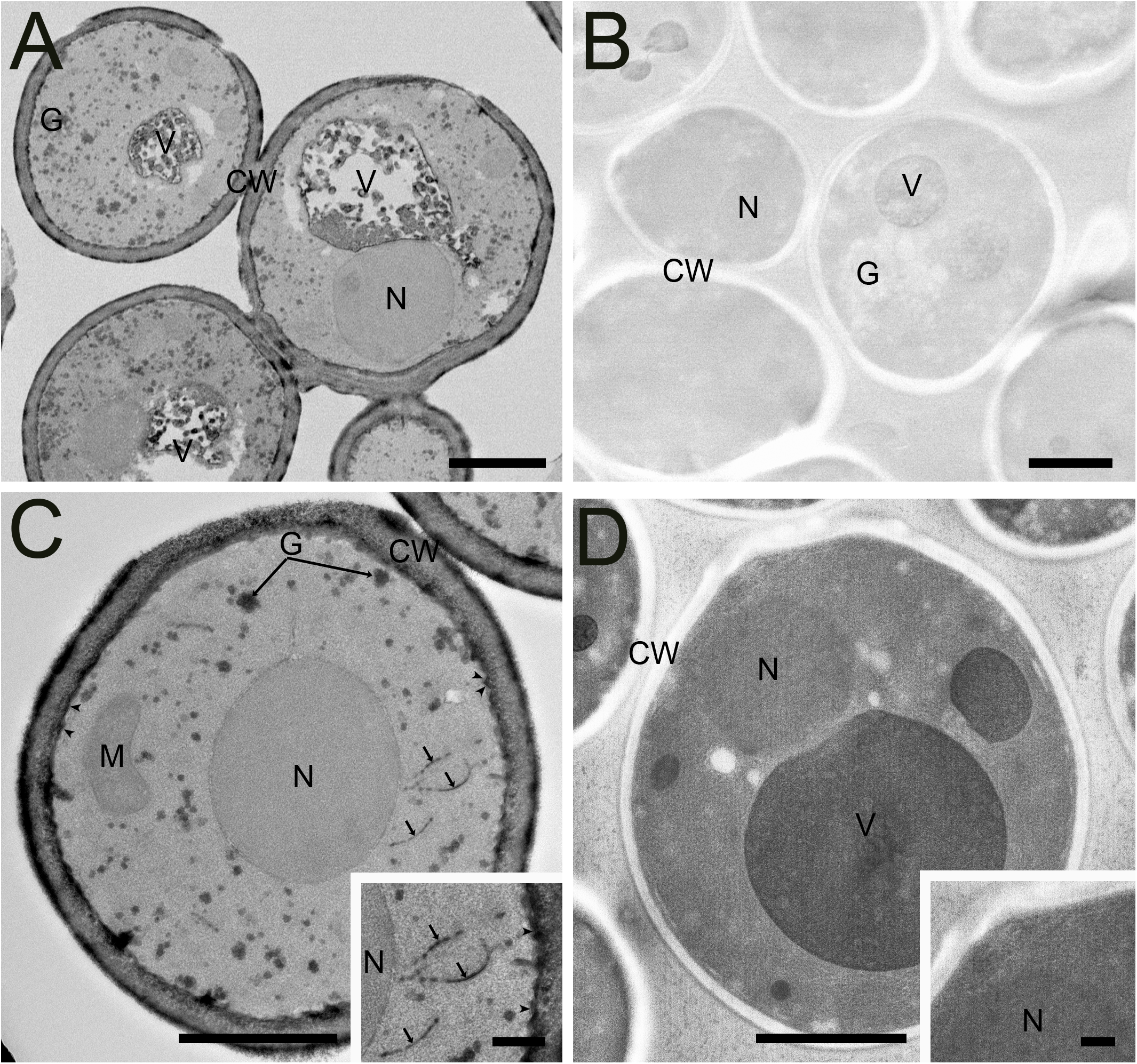
Comparison of high-pressure frozen yeast *S. cerevisiae* prepared by FSaqOTO and traditional AFS protocols and imaged via STEM and TEM. **A**. Overview STEM image of *S. cerevisiae* demonstrated very high contrast staining of the cell wall (CW), vacuolar contents and their delimiting membrane and glycogen using FAaqOTO and using identical imaging condition for **B** compared to a traditional AFS staining protocol. **C and inset**. FSaqOTO prepped yeast sample at increased magnification with well-labeled cell walls (CW), glycogen (G), plasmamembrane (arrowheads) cisternae (arrows), and elevated nuclear (N) and mitochondria (M) staining compared to AFS (**5D**) revealed. Scale = 1 µm. **D and inset**. Corresponding high magnification (compare to **5C**) with <u>identical</u> acquisition conditions of an AFS processed yeast <u>but</u> with a rescaled histogram demonstrated the normal complement of organelles but with much reduced contrast, especially cell membranes. A-D Scale = 1 µm, C and D insets = 100 nm.

## Discussion

Over the last decade, a number of studies have applied the benefits of improved structural preservation, realized by combining cryo-preservation and freeze-substitution with vEM techniques, including yeast (Wei *et al*., 2012), *C elegans* (Hall, Hartwieg and Nguyen, 2013; Rahman *et al*., 2021), plants (Czymmek *et al*., 2020; Reagan, B.B., Burch-SMith, 2022), *Drosophila melanogaster*, and mouse brain (Tsang *et al*., 2018), to name just a few. Continued improvements, such as by Guo et al. (2020), applied an organic solvent-based freeze-substitution protocol with a cocktail of osmium, tannic acid, uranyl acetate and potassium permanganate (without water) for improved membrane contrast and conductivity with FIB-SEM in a broad array of specimens (mouse brain, plants, algae, yeast and *C. elegans*). Notably, earlier investigations with freeze-substitution with TEM in yeast demonstrated the benefits of adding very small amounts of water to the substitution fluid (1-5%) to specifically enhance the visibility of some cell membranes that were otherwise poorly contrasted (Buser and Walther, 2008). Other studies also showed the benefits of water in the initial freeze-substitution fluid for preserving fluorescent protein signals (and other fluorophores) within acrylic resin blocks for both vEM, lending such samples to more efficient correlative microscopy workflows (Peddie *et al*., 2014). Furthermore, organic solvent freeze-substitution protocols (with water) followed by the transition to an aqueous buffer and Tokuyasu cryo-sectioning (Ripper, Schwarz and Stierhof, 2008) demonstrated notably improved antigenicity and morphology for several difficult plant, nematode and *Drosophila* specimens using the Tokuyasu protocol. A related strategy to combine the benefits of freeze-substitution followed by rehydration with correlative workflows, termed “CryoChem” was recently developed (Tsang *et al*., 2018). This versatile approach used 0.2% glutaraldehyde plus 5% water in acetone before transitioning to buffer before further processing for fluorescence microscopy (fluorescent protein or DRAQ5), diaminobenzidine (DAB) reaction for APEX2 labeling (Lam *et al*., 2015; Martell *et al*., 2017) followed by OTO-based heavy metalization, resin infiltration before X-ray microscopy and/or FIB-SEM.

Our vEM FSaqOTO protocol was differentiated from the other freeze-substitution aqueous rehydration protocols, as we did not have a requirement for non-osmicated tissues to initially preserve fluorescence or other chemistries. As such, we were able to obtain the full benefits of a strong initial fixation with 2% OsO_4_ (e.g., potentially reduced extraction) and a simplified protocol. Also of note, structures that are carbohydrate dense usually have very limited osmication and metal staining in standard EM protocols. However, in our work, we found that the addition of potassium ferrocyanide, followed by the direct exchange into OsO_4_ (without rinsing) per Hua (Hua, Laserstein and Helmstaedter, 2015), consistently provided very strong contrast of certain carbohydrate dense plant cell walls, yeast cell walls, the lumen of Golgi and secretory vesicles and glycogen (Fig. 2B-D, Fig. 3A & B, Fig. 5A & C). This observation is consistent with the well-known properties of potassium permanganate to enhance glycogen contrast in conventional fixation preparations (Revel, Napolitano and Fawcett, 1960). Not surprisingly, for plants, algae and yeast, when potassium permanganate was included in the freeze-substitution fluid of the FIBSEM protocol used by Guo (Guo *et al*., 2020), a similar increased labeling of cell walls in these organisms and glycogen in yeast was observed.

As mentioned previously, the addition of a small percentage of water to substitution fluid improves certain organelle membrane visibility in freeze-substitution (Buser and Walther, 2008). This is especially evident when we compare chloroplastic, thylakoid and grana membranes, which normally lack significant contrast in non-aqueous freeze-substitution preparations (Bourett, Czymmek and Howard, 1999; Bobik, Dunlap and Burch-Smith, 2014; Czymmek *et al*., 2020) but are very conspicuous in most conventional fixation (Kaneko and Walther, 1995; McDonald, 2014; Anderson *et al*., 2021). While the addition of water alone to the substitution will have some benefit, we observed that in combination with our FSaqOTO approach that chloroplast membranes were very prominent (Fig. 3C) with easy visualization of individual stacks of grana membrane and lumen via vEM will open doors to improved resolution, 3D visualization and quantification of these cryo-preserved structures in bulk samples. Furthermore, while not nearly as prominent an improvement as membranes in conventional fixation OTO methods, overall, our protocol did appear to enhance all cytoplasmic structures including cellular membranes of *C. elegans* and *S. cerevisiae* over our traditional AFS (compare Figs. 4A & C with 4B & D; Figs 5a & C with 5B and D). Indeed, much groundwork has been laid in the field for vEM of both *S. cerevisiae* (Winey *et al*., 1995; Buser and Walther, 2008; Wei *et al*., 2012; Wu *et al*., 2020) and *C. elegans* (Hall, Hartwieg and Nguyen, 2013; Rahman *et al*., 2020, 2021) with cryo-preservation and freeze-substitution. Specifically, our traditional AFS protocol used here was very similar substitution chemistry with prior Narayan lab work in *C. elegans*. However, here, virtually all structures (membrane or not) had elevated staining beyond our other standard method (Compare Fig. 4A & C with 4B & D, Fig. 5A & C with 5B & D) which benefited overall sample signal, contrast, and conductivity for vEM. We wanted to directly compare our FSaqOTO and traditional freeze-substitution AFS protocols using identical image acquisition and display conditions (Compare 4A with 4B, 5A with 5B) via STEM and TEM. Our comparison was specifically optimized for our FSaqOTO samples and then we reproduced the same conditions for the AFS processed samples. We note that for both our *C. elegans* and *S. cerevisiae* specimens, the beam conditions to acquire a high quality FSaqOTO image were inadequate for the AFS processed samples. While it is true that our acquisition conditions could be further optimized for our AFS prepared samples, our comparison sought to allow a side-by-side appreciation of the substantial sample staining improvement using the high-contrast FSaqOTO samples as the baseline. Furthermore, at higher magnification, the freeze-substitution staining of sensitive membrane structures (e.g. thylakoid, endoplasmic reticulum, nuclear envelope) could not be appreciably recovered in either specimen (Compare Fig. 4C with 4D, Fig. 5C with 5D), even with optimized imaging conditions. The metallization staining gains allowed in our tested samples also enabled the samples to tolerate increased beam dosage without noticeable beam damage artifacts. Indeed, elevation of the sample stain allowed enhanced conductivity, increased and more signal from selected sample features, which resulted in improved overall signal-to-noise and resolution. To appreciate this, comparison with previous work using a polar solvent-based OTO freeze-substitution protocol on *A. thaliana* anthers (Czymmek *et al*., 2020) and using the same SBF-SEM system with FCC, reflected the electron dosage this sample could tolerate (accelerating voltage 2.5 kV, beam current 1 pA, dwell-time 0.8 µs, z-slice 70 nm, pixel size 5 nm pixels). While for the barley anthers analyzed here, we used the same pixel size (5 nm) and probe current (1 pA), but were able to double the accelerating voltage to 5 kV with a 3.75-fold greater dwell-time (3 µs/pixel) as well as reliably cutting at a thinner z-slice of 50 nm (Figs 2 and 3, Supplementary Data Video 1). Likewise, our FSaqOTO prepared yeast 3D FIB-SEM volumes (Supplementary Data Video 2) versus our previous work using traditional freeze-substitution via FIB-SEM (Wei *et al*., 2012) had nearly identical beam acquisition conditions for our FSaqOTO samples (accelerating voltage 1.5 keV, probe current ∼1 nA). However, at 381 µsec/pixel average total dwell-time/image calculated from the earlier work versus FSaqOTO at 3 µsec/pixel average total dwell-time/image reflects over a 100-fold reduction in average pixel dwell-time/image. Thus, in both the anther and yeast instances, much improved throughput and/or signal-to-noise and could allow significant gains for many other types of similarly cryo-prepared vEM samples.

Not unlike the morphological differences in staining selectivity between conventional fixation OTO versus traditional non-OTO EM protocols, our FSaqOTO protocol had a different appearance and staining pattern when compared to traditional freeze-substitution. Notably, while the overall organelle/membrane enhancement appeared to be improved, the selectivity was less prominent compared to conventional fixation OTO protocols. Strictly aqueous-based conventional fixation OTO protocols have remarkably enhanced membrane contrast (Deerinck *et al*., 2018; Lippens *et al*., 2019). While FSaqOTO had a different appearance from other standard freeze-substitution protocols, it was characterized by consistently elevated tissue staining in all tested samples and fixation conditions. Furthermore, we noted that the cytoplasm and nucleoplasm of all cell types exhibited a granular/textured appearance that may reflect some level of nano-aggregation artifact of unknown origin. While this granular phenomenon may in part aid in the improved overall sample metallization and conductivity, it brings attention to the limits of the approach for certain questions that require high-fidelity, high-resolution ultrastructural studies.

Overall, our improved FSaqOTO method allowed more reliable vEM data acquisition and extended the experimental possibilities for cryo-preserved samples that are otherwise limited by low contrast and low conductivity organic solvent-based staining protocols. The addition of water in the substitution fluid and a straightforward solvent to water transition for heavy metal staining enabled noteworthy improvement in contrast with many structures. Furthermore, these samples realized a much-improved sample tolerance to electron beam dosage which enabled longer integration times and increased resolution resulting in better signal-to-noise for a given pixel size. Alternatively, the approach can allow increased throughput for freeze-substituted vEM studies enabling more samples or larger volumes per unit time. Finally, there are numerous opportunities for the modification and/or improvement of our staining strategy via adjustments to the substitution fluid (Ripper, Schwarz and Stierhof, 2008; Guo *et al*., 2020), aqueous OTO and metalization steps (Deerinck *et al*., 2018; Genoud *et al*., 2018) and/or combination with correlative approaches (Caplan *et al*., 2011; Tsang *et al*., 2018; Duncan *et al*., 2021). Ultimately, we hope that our modified freeze-substitution vEM staining strategy will stimulate more labs to explore other recipes for improved selection and enhancement of target cell structures while maintaining the full benefits of cryo-preservation and freeze-substitution.

## ACKNOWLEDGMENTS

We thank Mohammad Rahman and Orna Cohen-Fix, NIDDK, for the *C*.*elegans* samples, and Lucyna Lubkowska and Mikhail Kashlev, NCI, for the yeast samples. We acknowledge the Advanced Bioimaging Laboratory (RRID:SCR_018951) at the Donald Danforth Plant Science Center for support with sample preparation including screening samples with LEO 912AB TEM acquired through an NSF Major Research Instrumentation grant (DBI-0116650). We acknowledge the DOE BER (DBI-0116650) to K. Czymmek and USDA NIFA Grant No.2019-67013-29010 to B. C. Meyers. We thank Dr. Steven Goodman and Microscopy Innovations, Inc. for guidance on handling mPrep/s capsules under cryo-genic temperatures. We also thank Joel Mancuso and Ruth Redman (Zeiss) for excellent support and Lisa Chan (Zeiss) collecting the high-resolution SBF-SEM datasets of barley anther and root samples. This project has been funded in whole or in part with Federal funds from the National Cancer Institute, National Institutes of Health, under Contract No. 75N91019D00024. The content of this publication does not necessarily reflect the views or policies of the Department of Health and Human Services, nor does mention of trade names, commercial products, or organizations imply endorsement by the U.S. Government.

## AUTHOR CONTRIBUTIONS

KC conceived of approach, SB, HB, VB, KC, KD, and KN performed sample preparation and/or imaging. KC and KN wrote and edited manuscript. SB, KD, BCM edited manuscript.

