## Supplementary Data 1 for "A Versatile Enhanced Freeze-Substitution Protocol for Volume Electron Microscopy"

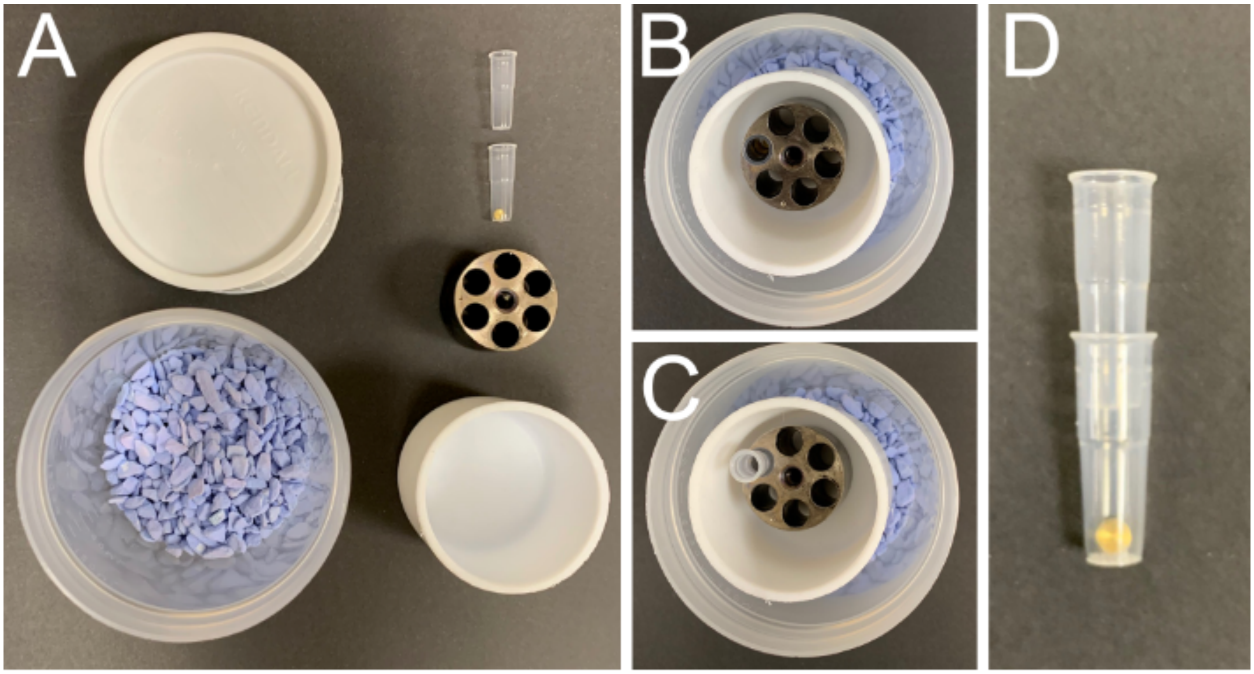


**Supplementary Data 1.** FSaqOTO mPrep capsules and containers used for freeze-substitution of plant materials at room temperature and shown here without reagents. **A.** Top view (start upper left and follow counterclockwise) of specimen cup lid, specimen cup with Drierite™ desiccant, Teflon™ substitution fluid container, mPrep capsule CPD holder, mPrep capsule bottom with HPF planchette (and specimen) inside, and the second mPrep/s capsule used as a top. **B.** Top view of assembled specimen cup with mPrep CPD holder within Teflon™ substitution fluid container in specimen cup. **C.** Same as B, but with a bottom mPrep/s capsule to hold HPF planchette for processing. **D.** Configuration of final mPrep capsule assembly to entrap the sample with HPF planchette (and specimen) between two mPrep/s capsules.
