## Supplementary Data 4 for "A Versatile Enhanced Freeze-Substitution Protocol for Volume Electron Microscopy"

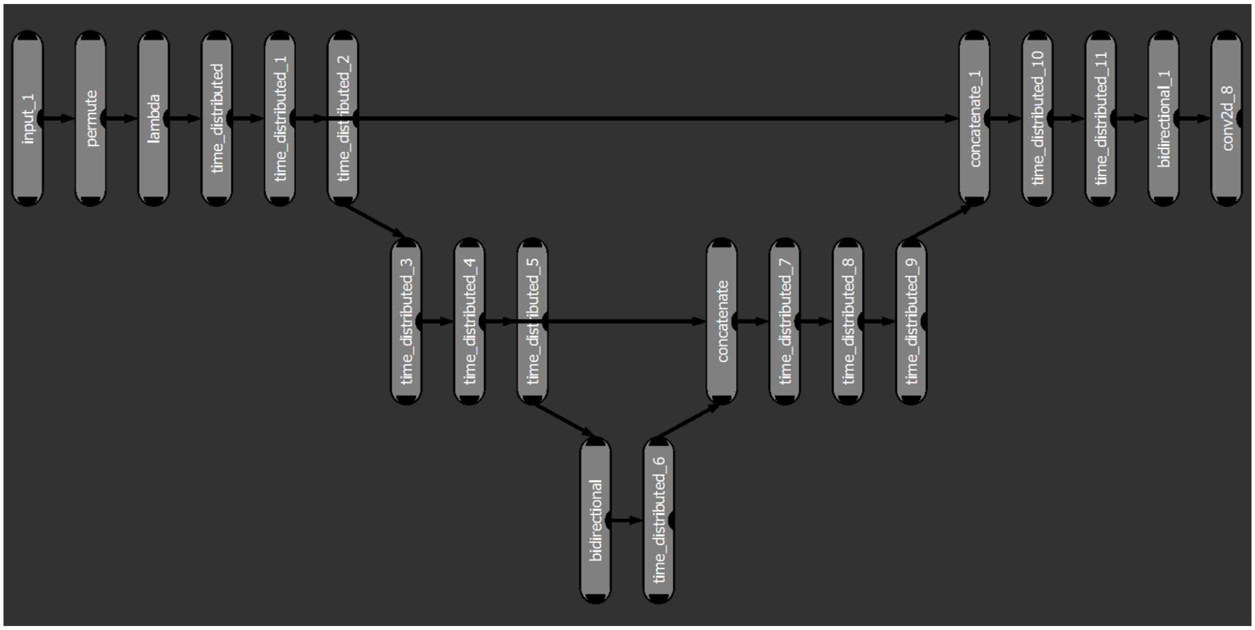


**Supplementary Data 4.** 3D Sensor Deep learning model parameters used with ORS Dragonfly segmentation of barley root (**Fig 2E**).
